# Retrieval of an ethanol-conditioned taste aversion promotes GABAergic plasticity in the insular cortex

**DOI:** 10.1101/2024.03.20.585950

**Authors:** Lisa R. Taxier, Meghan E Flanigan, Harold L. Haun, Thomas L. Kash

## Abstract

Blunted sensitivity to ethanol’s aversive effects can increase motivation to consume ethanol; yet, the neurobiological circuits responsible for encoding these aversive properties are not fully understood. Plasticity in cells projecting from the insular cortex (IC) to the basolateral amygdala (BLA) is critical for taste aversion learning and retrieval, suggesting this circuit’s potential involvement in modulating the aversive properties of ethanol. Here, we tested the hypothesis that GABAergic activity onto IC-BLA projections would be facilitated following the retrieval of an ethanol-conditioned taste aversion (CTA). Consistent with this hypothesis, frequency of mIPSCs was increased following retrieval of an ethanol-CTA across cell layers in IC-BLA projection neurons. This increase in GABAergic plasticity occurred in both a circuit-specific and learning-dependent manner. Additionally, local inhibitory inputs onto layer 2/3 IC-BLA projection neurons were greater in number and strength following ethanol-CTA. Finally, DREADD-mediated inhibition of IC parvalbumin-expressing cells blunted the retrieval of ethanol-CTA in male, but not female, mice. Collectively, this work implicates a circuit-specific and learning-dependent increase in GABAergic tone following retrieval of an ethanol-CTA, thereby advancing our understanding of how the aversive effects of ethanol are encoded in the brain.

**Significance statement:** Sensitivity to the aversive properties of ethanol contributes to motivation to consume alcohol. However, the plasticity-associated mechanisms through which ethanol’s aversive effects are represented within neural circuits are largely unidentified. In the present study, we used whole-cell patch clamp electrophysiology combined with synaptic input mapping to identify alterations in GABAergic plasticity within the insula, and within cells projecting from the insula to the basolateral amygdala. We demonstrate learning and circuit-specific alterations in GABAergic tone following retrieval of an ethanol-conditioned taste aversion, as well as a male-specific role for Parvalbumin-expressing interneurons in modulating the strength of an ethanol-conditioned taste aversion. Combined, these findings provide novel insights into how the aversive properties of ethanol are encoded within brain circuitry.

## Introduction

Alcohol has rewarding and aversive properties, both of which contribute to propensity to drink. Although the reinforcing properties of alcohol have received the lion’s share of attention, a growing body of literature highlights the fundamental role of alcohol’s aversive properties in motivation to drink. Higher sensitivity to the aversive effects of alcohol has been repeatedly linked to decreased propensity to consume alcohol (Mayfield et al., 2008; King et al., 2019), whereas lower sensitivity to the aversive effects of alcohol is associated with higher rates of alcohol use disorder (AUD) and binge drinking behaviors (Schuckit, 1994; King et al., 2011; Schuckit et al., 2014). The aversive properties of alcohol are frequently assessed in preclinical models by leveraging Pavlovian conditioned taste aversion (CTA) paradigms, in which rodents learn to avoid an innately rewarding tastant such as saccharin (the conditioned stimulus, or CS) after this tastant is paired with an intraperitoneal injection of ethanol (the unconditioned stimulus, or US), which induces malaise. Robust learned avoidance of the CS can be induced with a single CS-US pairing, allowing for precise control over timing and dose of ethanol exposure. Rodent models of the aversive effects of alcohol, leveraging ethanol-CTA, mirror clinical findings; mice or rats with higher sensitivity to alcohol’s aversive effects exhibit decreased propensity to drink alcohol, whereas rodents bred selectively for high drinking show diminished sensitivity to ethanol-CTA (Barkley-Levenson et al., 2015; Crabbe et al., 2019; Savarese et al., 2020).

The insular cortex (IC) represents a promising locus for encoding the aversive properties of alcohol. The IC is a densely interconnected structure, interfacing with several other regions to subserve a wide array of functions (Gogolla, 2017). Patients with AUD exhibit altered IC functional connectivity (Sullivan et al., 2013; Müller-Oehring et al., 2015; Manuweera et al., 2022), as well as altered IC activity in response to alcohol-associated cues (Zeng et al., 2021; Bach et al., 2024). In rodents, the IC is similarly vulnerable to alcohol, exhibiting structural and functional changes following chronic alcohol exposure (Chen et al., 2015; Marino et al., 2021). The IC is also an emergent player in modulating aversion-resistant drinking. For instance, digestion of perineuronal nets, extracellular matrix structures that surround Parvalbumin-expressing interneurons, within the IC reduced aversion-resistant drinking in mice (Chen and Lasek, 2020). Moreover, inhibiting glutamatergic inputs from the insula to the nucleus accumbens core blunted aversion-resistant drinking in rats (Seif et al., 2013). Importantly, the insula is directly responsive to ethanol-CTA. For example, cfos expression was associated with less saccharin consumption during ethanol-CTA retrieval in rats (Saalfield and Spear, 2019), suggesting that heightened aversion corresponds to greater activation of the IC. Additionally, levels of β-galactosidase, used as an index of neuronal activation, were elevated in the IC of adult rats 1 hour following ethanol-CTA retrieval (Gore-Langton et al., 2022). Combined, these data suggest that the IC encodes information regarding the aversive properties of ethanol; however, the plasticity-associated mechanisms within the IC responsible for modulating ethanol aversion-associated behaviors remain largely uncharacterized.

Although the mechanisms through which the aversive effects of ethanol may be represented in the IC are unclear, work exploring basic mechanisms of other taste aversion memories may lend insight. Importantly, protein synthesis and gene expression alterations within the IC are required for a lithium chloride (LiCL)-induced CTA memory across multiple memory phases (Rosenblum et al., 1993; Hadamitzky et al., 2016). Reciprocal connectivity between the IC and the basolateral amygdala (BLA) has also been repeatedly implicated in taste aversion memory (Buresová, 1978; Kayyal et al., 2019; Abe et al., 2020). Lastly, taste learning via retrieval of a LiCl-conditioned taste aversion promoted GABAergic plasticity onto cells projecting from the IC to the BLA (Yiannakas et al., 2021). This existing literature suggests that inhibitory plasticity within IC-BLA circuitry is critical for CTA learning; however, whether similar mechanisms are at play for ethanol-CTA learning is unknown.

Despite extensive evidence linking lower sensitivity to the aversive effects of alcohol to higher incidence of AUD, how the aversive properties of alcohol are functionally encoded within brain circuitry remains an open question. Previous work demonstrating the importance of GABAergic plasticity in an IC-BLA circuit in tase aversion learning led us to hypothesize that this circuitry is also critically involved in retrieval of an ethanol-CTA. Therefore, the goal of the present study was to evaluate whether ethanol-CTA retrieval produces GABAergic plasticity in IC-BLA circuitry. We also employed local synaptic input mapping to assess whether cells projecting from the IC to the BLA receive modified local inhibition following ethanol-CTA retrieval. Lastly, we inhibited parvalbumin (PV)-expressing interneurons within the IC to determine the necessity of these cells for retrieval of an ethanol-CTA. Our results suggest that retrieval of an ethanol-CTA promotes GABAergic plasticity onto IC-BLA projection neurons in a learning-dependent fashion, and that local inhibitory input onto IC-BLA projection neurons is altered in a layer-specific manner following ethanol CTA. Moreover, PV interneuron activity, in males only, modulates the strength of an ethanol CTA. Combined, these findings implicate plasticity of an IC-BLA circuit as a key player in encoding the aversive properties of ethanol, and suggest that local inhibitory plasticity onto IC-BLA projecting cells occurs as a result of ethanol CTA retrieval.

## Materials and methods

### Animals

Male and female C57Bl/6 mice (n=52; Stock #: 000664, Jackson Laboratories) were obtained from Jackson Laboratories at 8 weeks of age. Parvalbumin-Cre (PV-Cre, n=36) mice were obtained from our in-house breeding colony. Mice were maintained on a 12-hour reverse light-dark cycle in temperature and humidity-controlled facilities, and were given ad libitum access to food (Prolab Isopro RMH 3000, LabDiet) and water. Mice were group-housed until surgery, and were subsequently single-housed to allow for accurate monitoring of fluid consumption. Following surgery, which was conducted three weeks prior to the start of behavioral testing, mice remained singly housed for the duration of the experiment. All procedures followed the National Institutes of Health Guide for the Care and Use of Laboratory Animals and were approved by the University of North Carolina-Chapel Hill School of Medicine Institutional Animal Care and Use Committee.

### Surgical procedures

Following one week of acclimation to the animal facility, mice underwent a single surgical procedure during which they were bilaterally infused into the BLA (-.9AP, +/-3.45 ML, -4.95 DV) with .2 µL AAV-rg-CAG-tdTomato to label IC-BLA projection neurons for slice electrophysiology recordings and synaptic input mapping experiments. For DREADD experiments, PV-Cre mice were bilaterally infused into the insula (1.6AP +/-3.1ML, -3.4 DV) with .2 µL AAV8-hSyn-DIO-mCherry or AAV8-hSyn-DIO-hM4D(Gi)-mCherry. Mice were provided a single dose of subcutaneous Ketoprofen (5mg/kg) at the start of surgery for analgesia, and one dose per day for two days following surgery for postoperative analgesia, or until presurgical body weight was recovered. Mice recovered for three weeks prior to the start of behavioral experiments to allow for viral expression.

### Ethanol-conditioned taste aversion

Ethanol-CTA was conducted across three days, with each daily session separated by 24 hours. All sessions were conducted in the home cage. Mice were habituated to intraperitoneal (i.p.) injections for three days prior to the start of habituation. On day 1 (habituation), mice were given 30 minutes of access to a 0.5% saccharin solution. On day 2 (conditioning), mice were again given 30 minutes of access to a 0.5% saccharin solution, immediately followed by an i.p. injection of vehicle (saline) or 2g/kg EtOH. On day 3 (retrieval), mice were again given 30 minutes of access to 0.5% saccharin. For DREADD experiments, ethanol-CTA was conducted as described, with the addition of a .1mg/kg i.p. injection (Nagai et al., 2020) of the DREADD actuator Deschloroclozapine dihydrochloride (DCZ, Hello Bio) given 30 minutes prior to saccharin access on retrieval day. Bottles were weighed prior to and after each saccharin presentation session.

### Slice electrophysiology

Whole-cell patch-clamp recordings were obtained from the IC. Following rapid decapitation, brains were extracted and immersed in a chilled sucrose cutting solution (194mM sucrose, 20mM NaCl, 4.4mM KCl, 2mM CaCl_2_, 1mM MgCl_2_, 1.2mM NaH_2_PO_4_, 10mM D-glucose, and 26mM NaHCO) oxygenated with 95% O_2_/5% CO_2_. Coronal slices (250 uM) containing the IC were prepared on a vibratome (VT1200S, Leica) and transferred to oxygenated aCSF (124 mM NaCl, 4.4 mM KCl, 1 mM NaH2PO4, 1.2 mM MgSO4, 10 mM D-glucose, 2 mM CaCl2, and 26 mM NaHCO3) maintained at 35°C for a 30 min recovery period prior to recording. A cesium chloride-based internal solution (130 mM CsCl, 1 mM EGTA, 10 mM HEPES, 2 mM Mg-ATP, .2 mM Na2-ATP) was used during recordings of miniature inhibitory post-synaptic currents (mIPSCs), and a potassium gluconate-based internal solution (135 mM K-gluconate, 5 mM NaCl, 2 mM MgCl2, 10 mM HEPES, 0.6 mM EGTA, 4 mM Na2ATP, 0.4 mM Na2GTP) was used during recordings of resting membrane potential (RMP) during tests of DCZ functionality. To pharmacologically isolate mIPSCs, tetrodotoxin (TTX, Tocris, 500nM), CNQX (abcam, 20 µM), and D-AP-5 (abcam, 50µM) were added to the circulating aCSF. Recording pipettes (2–4 MΩ) were pulled from thin-walled borosilicate glass capillaries using a P95 pipette puller (Sutter Instruments). Signals were acquired using an Axon Multiclamp 700B amplifier (Molecular Devices), digitized at 10 kHz, filtered at 3 kHz, and analyzed in pClamp 10.7 (RMP) or Easy Electrophysiology (mIPSCs). Series resistance (*R*_s_) was monitored without compensation, and changes in *R*_s_ exceeding 20% were used as exclusion criteria.

### Synaptic input mapping

Slices were prepared as described above. Synaptic input mapping was conducted with Laser Assisted Stimulation and Uncaging software (LASU, Scientifica). RuBi-glutamate (300 µM) was supplied in 35 mL recirculating ACSF and photolysed using a pulsed 405 nm laser beam (25 mw) at 5x magnification across a stimulus grid (200 uM spacing) encompassing the entirety of the IC, including all cortical layers. Uncaging at each stimulation site was guided by mirror galvanometers controlled by LASU software, and was separated by 5s intervals. To isolate IPSCs, maps were recorded at a holding potential of +10 mV and a cesium methanesulfonate internal solution (135 mM cesium methanesulfonate, 10 mM KCl, 1 mM MgCl2, 0.2 mM EGTA, 4 mM MgATP, 0.3 mM Na2GTP, 20 mM phosphocreatine, with 1mg/mL QX-314) was used. IPSCs were observed within a 2–50 ms window after the stimulus for each sweep (Brill et al., 2016). Amplitude of each uncaging-evoked event was recorded in pClamp 10.7. Individual synaptic input maps were combined to create averaged synaptic input maps for both layer 2/3 and layer 5/6 IC cells by recording the amplitude, in pA, and x and y distance, in microns, of each responsive stimulation site from the soma. These values were then binned at 100 uM intervals. Smooth contours were derived in MATLAB using linear interpolation between 100 μm bins.

### Histology

To verify accuracy of viral targeting of DREADD constructs, mice underwent transcardial perfusion with ice-cold phosphate-buffered saline followed by 4% paraformaldehyde. Coronal slices containing the IC (30 µM) were collected using a vibratome (VT1200, Leica). Sections were mounted on SuperFrost plus slides and coverslipped with mounting media containing DAPI. Images were collected using a Keyence BZ-X800 microscope at 10x magnification. Mice with off-target expression (n=5), including one animal intended for functional electrophysiological validation of DREADD function, were excluded from analysis.

### Statistical Analyses

Statistical analyses were performed using GraphPad Prism 10 software (La Jolla, Ca). Behavioral data were analyzed using two-way ANOVAs using sex (male vs female) and treatment (saline vs ethanol) as dependent variables. Two-way ANOVAs using sex (male vs female) and treatment (saline vs ethanol) as dependent variables were used to compare between-group differences in electrophysiological measures. Two-way ANOVAs using treatment (saline vs ethanol) and distance to soma as dependent variables were used for synaptic input mapping experiments to compare summed cumulative synaptic inputs, and unpaired t-tests were used to compare total number of synaptic input sites in both layer 2/3 and layer 5/6. A two-way ANOVA using sex (male vs female) and treatment (saline vs ethanol) was used to assess DREADD-mediated differences in CTA learning. Significant ANOVA interactions were followed by Tukey’s *post hoc* comparisons, with the exception of significant interactions for summed cumulative synaptic input across the intra-or interlaminar extent, which were followed by Šídák’s multiple comparisons test, with significance set at *p* < 0.05.

## Results

### mIPSC frequency, but not amplitude, is higher in cells projecting from the IC to the BLA following ethanol-CTA retrieval

We first endeavored to determine the extent to which retrieval of an ethanol-CTA alters GABAergic plasticity onto cells projecting from the IC to the BLA. We hypothesized that ethanol-CTA retrieval would result in heightened GABAergic tone onto IC-BLA projecting cells. To test this hypothesis, we injected mice bilaterally into the BLA with a retrograde AAV construct to label IC-BLA projecting pyramidal cells (Fig 1B). Three weeks post-surgery, mice underwent ethanol-CTA (Fig 1A). We then obtained whole-cell patch clamp recordings from layer 2/3 and layer 5/6 IC-BLA projection neurons 1 hr following ethanol-CTA retrieval. A two-way ANOVA revealed a significant effect of treatment, such that both male and female mice exhibited a robust aversion to saccharin 24 hrs following a 2g/kg conditioning dose of EtOH relative to saline-conditioned controls (Fig 1C; *F*_(1,20)_ = 36.13, *p* < 0.0001). Consistent with our hypothesis, IC-BLA cells from mice that underwent ethanol-CTA had a higher frequency of miniature inhibitory post synaptic currents (mIPSCs) compared to cells from saline-conditioned mice, although amplitude remained unchanged. Higher frequency of mIPSCs was observed in both IC-BLA projections from layer 2/3 (Fig 2A-B; *F*_(1,29)_ = 8.951, *p* = 0.006), and from layer 5/6 (Fig 2D-E; *F*_(1,24)_ = 11.71, *p* = 0.002). No effects of ethanol-CTA retrieval were observed in Td-Tomato-negative cells (layer 2/3, Fig 2G-I; layer 5/6, Fig 2K-L), suggesting that altered GABAergic plasticity following ethanol-CTA was limited to IC-BLA projections.

**Figure 1.**
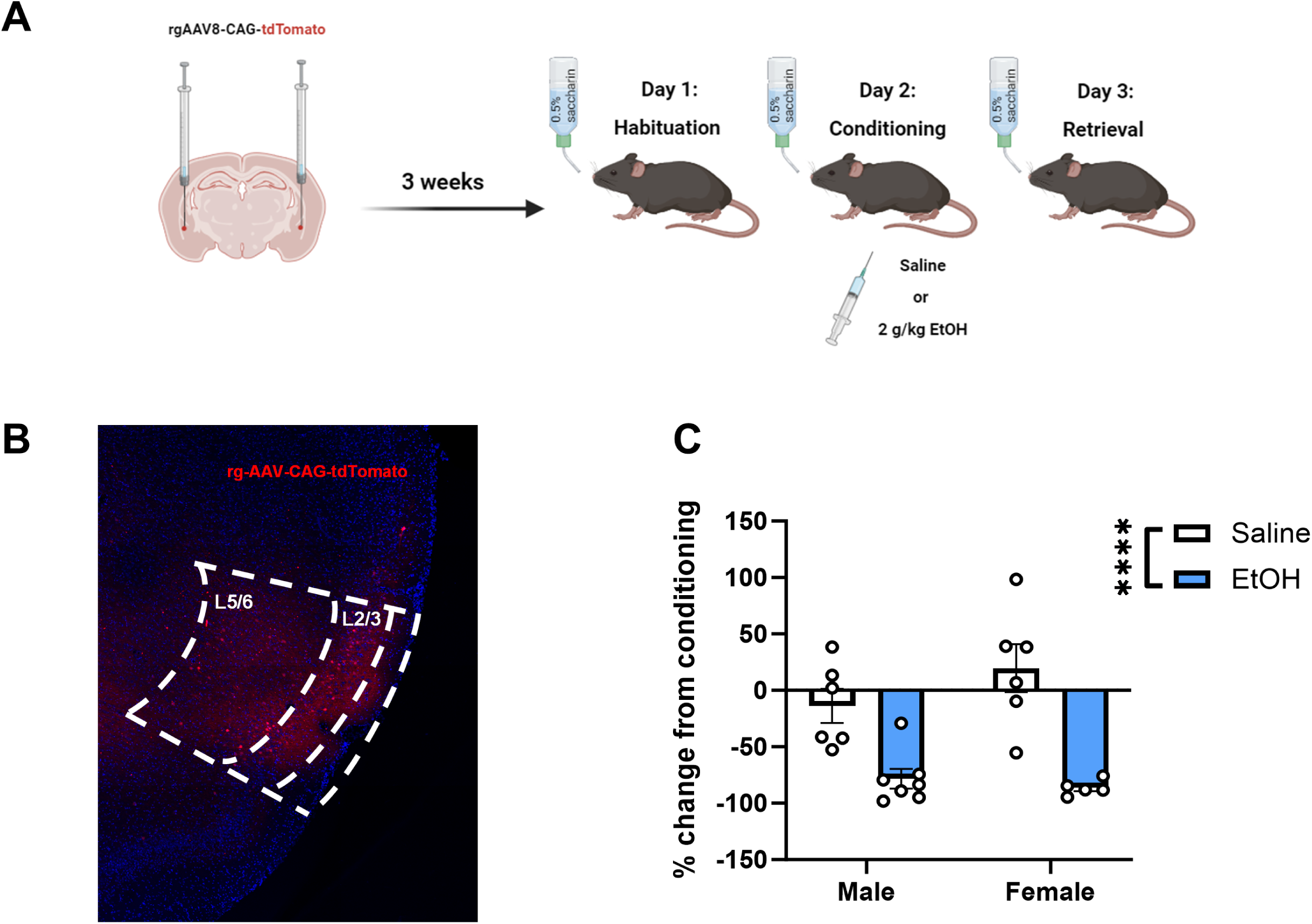
Male and female mice exhibit a robust conditioned taste aversion to a 2 g/kg dose of ethanol. (A) Experimental design. Mice received bilateral intra-basolateral amygdala (BLA) injections of a retrograde AAV construct allowing for expression of fluorescent tdTomato in cells projecting from the anterior insula (IC) to the BLA prior to ethanol-CTA. (B) A representative image of tdTomato expression indicating that viral expression was largely confined to the IC and present throughout the intralaminar extent. (C) Both male and female mice receiving a 2 g/kg i.p. injection of ethanol drank significantly less saccharin during retrieval compared to saline-conditioned controls (\*\*\*\**p* < 0.0001 main effect of ethanol, data expressed as % change from conditioning).

**Figure 2.**
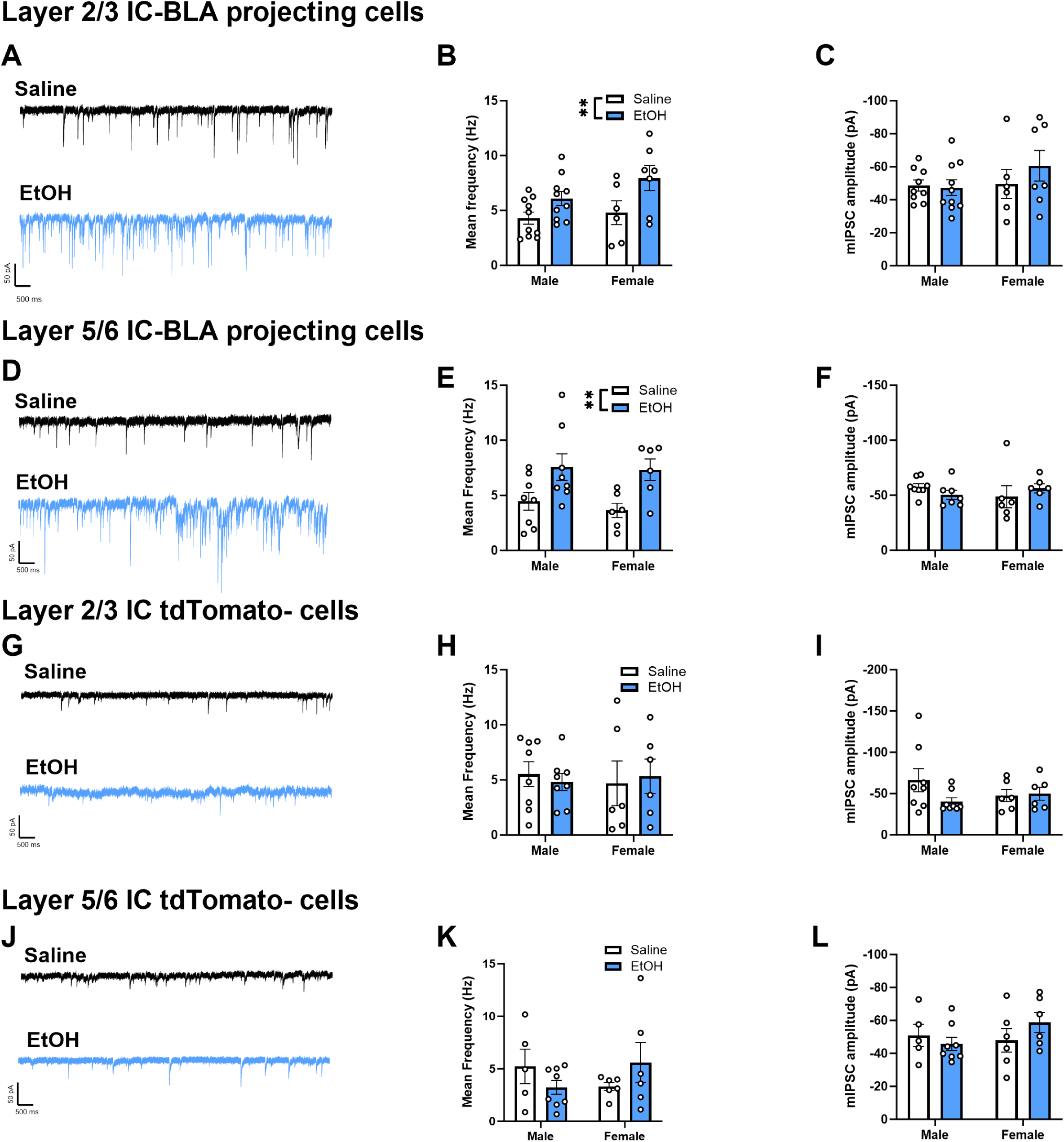
mIPSC frequency, but not amplitude, is higher in cells projecting from the IC to the BLA following ethanol-CTA retrieval. Representative traces of mIPSCs from layer 2/3 (A) and layer 5/6 (D) IC-BLA projecting neurons 1 hr following saline or ethanol-CTA retrieval. Frequency of mIPSCs was significantly higher in cells from ethanol-conditioned mice compared to saline-conditioned controls, in both layer 2/3 (B) and layer 5/6 (E), whereas amplitude was unaltered (C, F). Representative traces of mIPSCs from the general population (tdTomato-) of IC pyramidal cells in layer 2/3 (G) and layer 5/6 (J) 1 hr following saline or ethanol-CTA retrieval. Both frequency (H,K) and amplitude (I,L) of mIPSCs in tdTomato-cells were unchanged by retrieval of ethanol-CTA compared to saline-conditioned controls, regardless of layer (\*\**p*<0.01, main effect of ethanol).

### A single dose of ethanol does not elicit ethanol-CTA retrieval-induced changes in GABAergic plasticity

In order to determine whether the effects of ethanol-CTA retrieval on GABAergic plasticity were dependent upon retrieval of ethanol-CTA versus ethanol alone, we next administered a single i.p. dose of 2g/kg ethanol, and obtained whole-cell patch clamp recordings from layer 2/3 and layer 5/6 IC-BLA projecting cells 24 hrs later. There were no significant differences in mIPSC frequency or amplitude in IC-BLA projecting cells 24 hrs following i.p. injection of EtOH in the absence of CTA learning, regardless of layer (layer 2/3, Fig 3A-C; layer 5/6, Fig 3D-F). These data suggest that the effects of EtOH on GABAergic plasticity following CTA retrieval are learning dependent.

**Figure 3.**
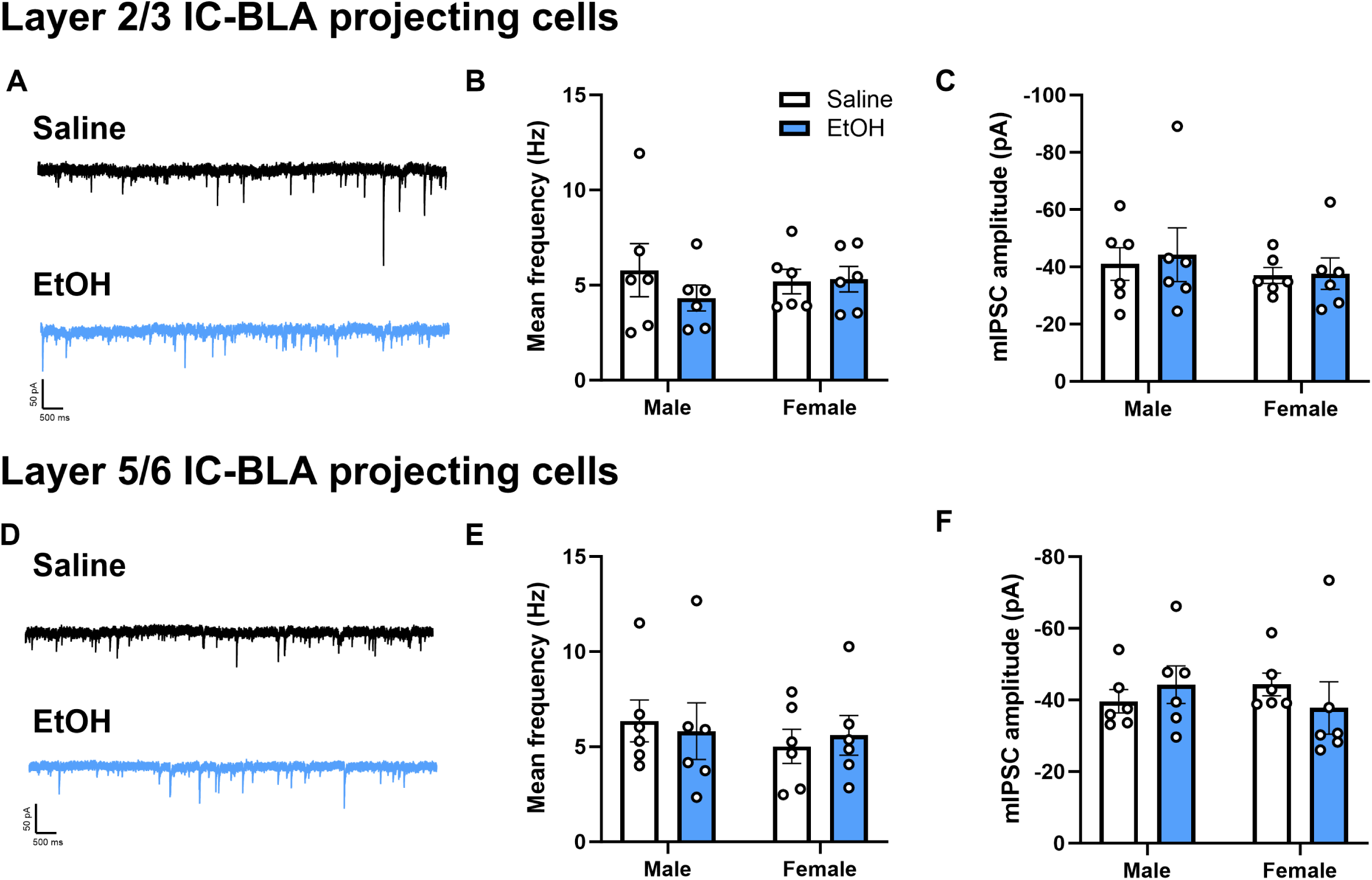
The effects of ethanol-CTA retrieval on GABAergic plasticity within an IC-BLA circuit are learning-dependent. Representative traces of mIPSCs from layer 2/3 (A) and layer 5/6 (D) IC-BLA projecting neurons 24 hr following i.p. injection of ethanol. Both frequency (B, E) and amplitude (C,F) of mIPSCs in IC-BLA projection neurons were not significantly altered 24 hr after i.p. injection of ethanol.

### Local inhibitory input onto layer 2/3 IC-BLA projecting cells is greater following ethanol-CTA retrieval

Because local inhibition plays a critical role in modulating pyramidal cell excitability, and previous work implicates inhibitory plasticity within the IC as a key feature of taste aversion learning (Yiannakas et al., 2021), we next sought to determine whether ethanol-CTA retrieval produces alterations in local inhibitory inputs onto IC-BLA projecting cells. We hypothesized that there would be an increase in local inhibition onto IC-BLA projecting cells following ethanol-CTA retrieval. To test this hypothesis, we conducted synaptic input mapping experiments using laser-applied stimulation and uncaging (LASU) of the caged glutamate compound, Rubi-glutamate. Whole-cell patch clamp recordings were obtained from layer 2/3 and layer 5/6 IC-BLA projecting cells 1 hr following ethanol-CTA retrieval (Fig 4A). A point grid with 200µm spacing between points was overlaid across the entirety of the IC (Fig 4B), and focal laser stimulation was sequentially applied while recording uncaging-evoked IPSCs from the patched cell in order to determine the presence, strength, and spatial location of inhibitory inputs to recorded cells. Resulting input maps (Fig 4C) revealed stronger local inhibitory input onto layer 2/3 IC-BLA projecting cells following ethanol-CTA retrieval compared to cells from mice conditioned with saline. There were more total inhibitory synaptic input sites to layer 2/3 IC-BLA projecting cells in ethanol-CTA retrieving mice compared to saline-treated mice (Fig 4D; *t*_(14)_ = 2.34, *p* = 0.03). When cumulative amplitude was collapsed across the interlaminar extent (i.e., horizontal distance to soma), two-way ANOVAs revealed a main effect of dorsal/ventral distance (*F*_(17, 612)_ = 123.5, *p* < 0.0001), such that amplitude of inhibitory events was higher at uncaging sites nearer the soma, and treatment (*F*_(1,612)_ = 421.5, *p* < 0.0001), such that amplitude of uncaging-elicited events was higher in cells from ethanol-conditioned mice compared to saline-conditioned controls (Fig 4E). Two-way ANOVA also yielded a significant interaction between distance to soma and treatment along the dorsal/ventral extent (*F*_(17,612)_ = 40.64, *p* < 0.0001). The same effects were present when cumulative amplitude of uncaging-evoked events in layer 2/3 cells was collapsed across the intralaminar distance (i.e., vertical distance to soma); a two-way ANOVA revealed a significant effect of horizontal distance (*F*_(15, 544)_ = 127.3, *p* < 0.0001) such that amplitude of inhibitory events was higher when glutamate was uncaged at sites in close horizontal proximity to the recorded cell’s soma (Fig 4F). Additionally, the effect of treatment was significant (*F*_(1, 544)_ = 454.1, *p* < 0.0001) such that IPSC amplitude was greater along the horizontal plane in cells from ethanol-conditioned mice compared to cells from saline-conditioned mice. A two-way ANOVA also yielded a significant interaction between distance to soma and treatment along the medial/lateral plane (*F*_(15, 544)_ = 29.95, *p* < 0.0001). Combined, these data demonstrate that local inhibitory input onto layer 2/3 IC-BLA projecting cells is greater following ethanol-CTA retrieval, particularly from cells in close proximity to IC-BLA projecting neurons,

**Figure 4.**
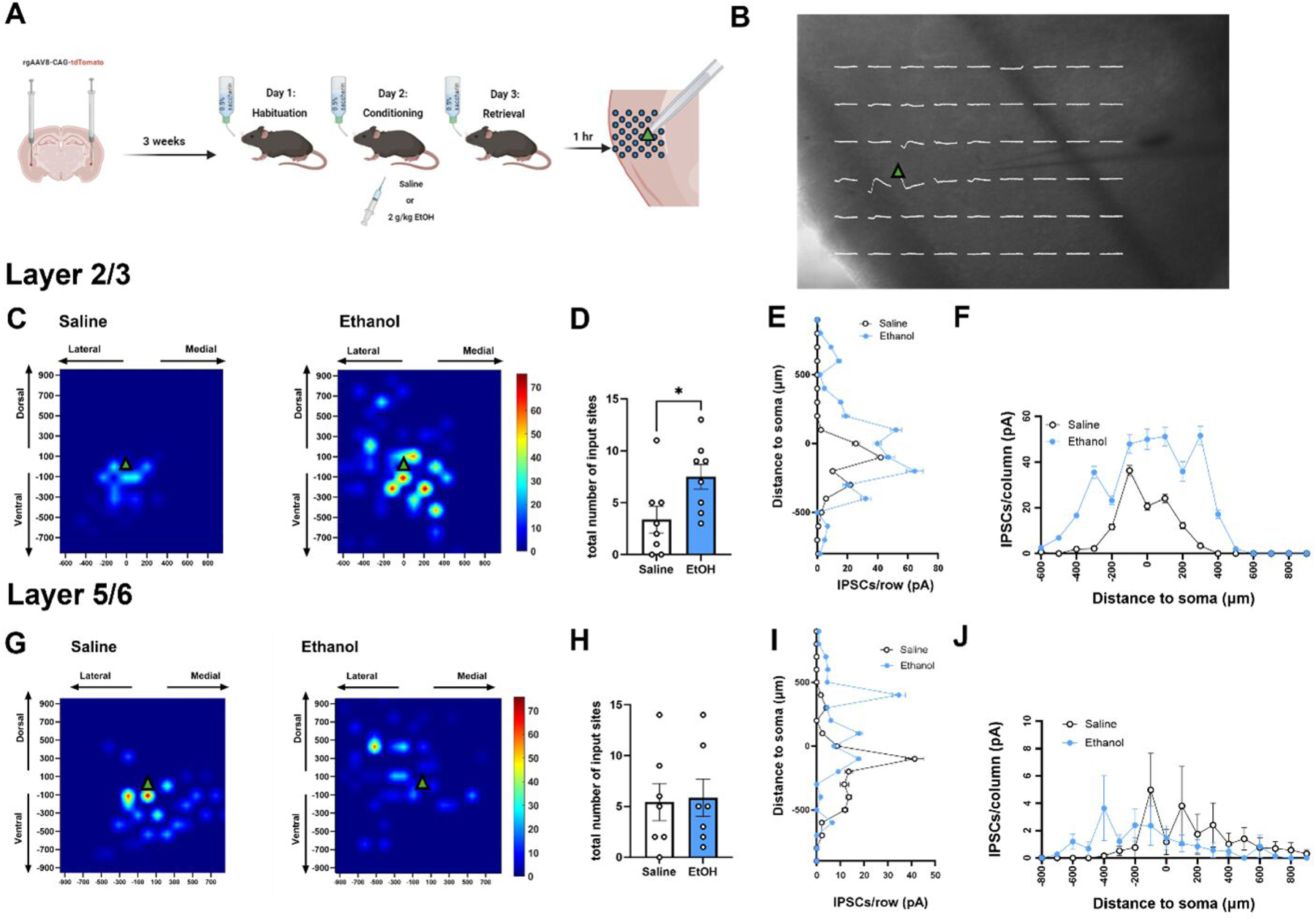
Local inhibitory input onto layer 2/3 IC-BLA projecting cells is greater following ethanol-CTA retrieval. (A) Experimental design. Mice received bilateral intra-basolateral amygdala (BLA) injections of a retrograde AAV construct allowing for expression of fluorescent tdTomato in cells projecting from the anterior insula (IC) to the BLA prior to ethanol-CTA. 1 hr following ethanol-CTA retrieval, synaptic input maps to IC-BLA projecting cells were created using laser applied stimulation and uncaging. (B) Representative spatially localized uncaging-elicited events (white traces) are overlaid on a coronal section containing the IC. Green triangle indicates approximate location of cell soma in (B), (C), and (G). (C) Averaged input maps from IC-BLA layer 2/3 cells from saline-conditioned (left) or ethanol-conditioned mice (right). The number of total inhibitory input sites to layer 2/3 cells was greater in ethanol-conditioned mice relative to saline-conditioned controls (D). In layer 2/3, cumulative amplitude of uncaging-evoked events in pA in increments of 100µm from the soma across interlaminar (E) and intralaminar (F) extents was higher in ethanol-conditioned mice relative to saline-conditioned controls, particularly at locations nearest the soma, as indicated by significant main effects of distance and treatment, and a distance x treatment interaction. (G) Averaged input maps from IC-BLA layer 5/6 cells from saline-conditioned (left) or ethanol-conditioned mice (right). The number of total inhibitory input sites to layer 5/6 cells was not significantly different in ethanol-conditioned mice relative to saline-conditioned controls (H). In layer 5/6, cumulative amplitude of uncaging-evoked events in pA in increments of 100µm from the soma across the interlaminar extent (I) was impacted by a significant effect of distance, and a distance x treatment interaction such that saline-conditioned mice received stronger inhibitory inputs from ventral locations relative to ethanol-conditioned mice, and ethanol-conditioned mice received stronger inhibitory inputs from dorsal locations relative to saline-conditioned mice. Although there was no main effect of treatment nor an interaction, there was a significant main effect of distance along the interlaminar extent suggesting increased amplitude of uncaging-evoked events nearer to the cell body. (\**p* < 0.05, significant effect of ethanol).

In contrast to layer 2/3, composite inhibitory input maps to Layer 5/6 IC-BLA projecting cells revealed reorganization of inhibitory synaptic inputs without significant differences in total number of input sites (Fig 4H). When cumulative amplitude was collapsed across the interlaminar extent, two-way ANOVA yielded a significant main effect of distance along the dorsal/ventral axis (*F*_(18,646)_ = 105.3, *p* < 0.0001), as well as a significant interaction between distance and treatment (*F*_(18,646)_ = 66.16, *p* < 0.0001). Sidak’s *post-hoc* comparisons indicated that saline-conditioned mice received stronger inhibitory inputs from more ventral locations compared to ethanol-conditioned mice, whereas ethanol-conditioned mice received stronger inhibitory inputs from more dorsal locations compared to saline-conditioned mice (Fig 4I). Although two-way ANOVA revealed no significant main effect of treatment when cumulative amplitude was collapsed across the intralaminar extent, nor an interaction, there was a significant main effect of distance along the medial/lateral axis (*F*_(17,612)_ = 1.719, *p* = 0.04; Fig 4J). Collectively, these data suggest that inhibitory input onto layer 5/6 IC-BLA projecting cells is reorganized along the dorsal-ventral axis following ethanol-CTA retrieval, but not in a manner consistent with increased total local inhibitory input.

### Inhibition of parvalbumin-expressing interneurons within the IC blunts the expression of ethanol-CTA in male mice

Previous work suggests that PV interneurons within the IC are critical for the expression of a LiCl CTA. This finding, coupled with our own finding that local inhibitory inputs to L2/3 IC-BLA projections are strengthened following ethanol-CTA retrieval, led us to hypothesize that IC PV interneurons are likewise key for ethanol-CTA retrieval. To test this hypothesis, we employed DREADD-mediated inhibition of PV interneurons within the IC of PV-Cre mice just prior to ethanol-CTA retrieval (Fig 5A). We used DCZ as an actuator for Gi-coupled DREADDs specifically expressed in IC PV interneurons (Fig 5B) 30 minutes prior to ethanol-CTA retrieval, and first validated that DCZ on its own did not significantly modulate CTA (data not shown). We also conducted electrophysiological validation of DCZ as an actuator of Gi-coupled DREADDs in PV interneurons. A two-way ANOVA revealed a significant interaction (*F*_(1,32)_ = 4.205, *p* = 0.05) between DREADD condition (mCherry vs Gi) and DCZ application, demonstrating that bath application of DCZ resulted in a significant hyperpolarization of membrane potential only in mice expressing Gi DREADDs (Fig 5C, *p* = 0.04). DREADD-mediated inhibition of PV interneurons within the IC resulted in a blunting of ethanol-CTA (Fig 5D); a two-way ANOVA revealed a significant effect of DREADD condition (mCherry vs Gi; *F*_(1,27)_ = 6.680, *p* = 0.016), as well as a significant interaction between sex and DREADD condition (*F*_(1,27)_ = 4.295, *p* = 0.048). A Tukey’s *post hoc* test demonstrated that Gi-DREADD mediated inhibition of PV interneurons blunted the expression of ethanol-CTA in male mice (*p* = 0.02), but not female mice. These data implicate PV interneurons in the male IC as important mediators of EtOH CTA retrieval.

**Figure 5.**
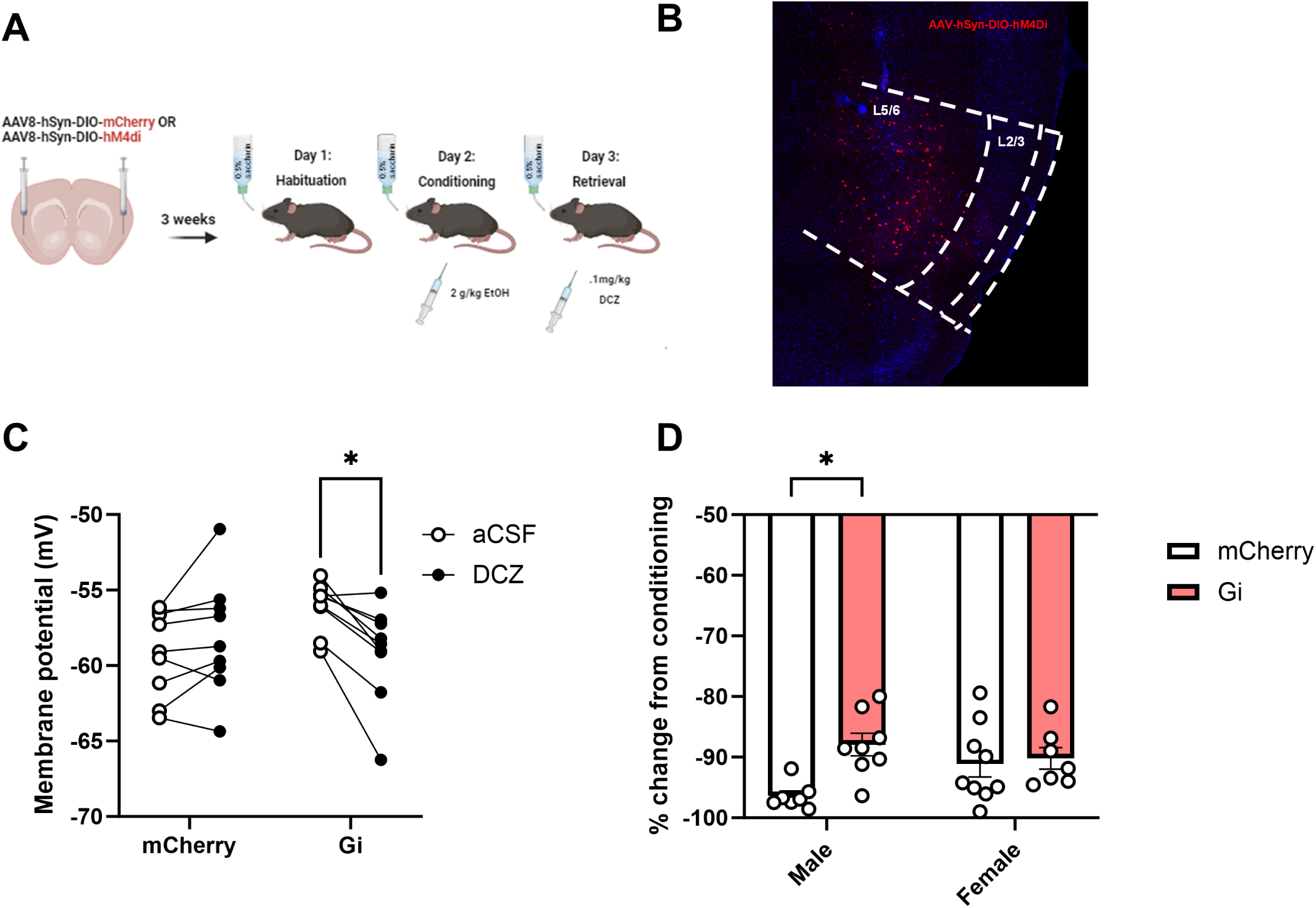
Inhibition of parvalbumin-expressing interneurons within the IC blunts the expression of ethanol-CTA in male mice. (A) Experimental design. PV-Cre mice received bilateral intra-insular injections of Cre-dependent mCherry or hM4di constructs, allowing for expression of either mCherry (controls) or Gi DREADDs three weeks prior to undergoing ethanol-CTA. Mice were given a .1mg/kg i.p. injection of DCZ 30 min prior to retrieval on Day 3. (B) Representative image of AAV-hSyn-DIO-hM4Gi confined largely within the IC. (C) Electrophysiological validation of DREADD function. Bath application of 300nM DCZ resulted in a significant hyperpolarization of membrane potential in Gi-expressing cells, but not mCherry-expressing cells. (D) A significant effect of DREADD condition and a significant interaction between sex and DREADD condition indicated that male, but not female mice expressing the hM4Gi construct drank significantly less saccharin during retrieval relative to mCherry-expressing controls following a.1 mg/kg i.p. injection of DCZ. (\**p* < 0.05 male mCherry vs male Gi, data expressed as % change from conditioning).

## Discussion

Converging evidence implicates the IC as a multimodal integration site that exhibits structural and functional alterations in response to aversive stimuli. Here, we show robust increases in GABAergic tone in local insula circuitry, as well as onto IC-BLA projecting cells, following ethanol-CTA retrieval, suggesting a novel role for GABAergic plasticity in IC circuitry in encoding the aversive effects of ethanol. Furthermore, we demonstrate that increased frequency of mIPSCs onto IC-BLA projecting cells following ethanol-CTA occurs in a learning-dependent manner, consistent with a role for GABAergic plasticity in IC-BLA circuitry in retrieval of an aversive taste memory. Lastly, we show that inhibiting activity of PV interneurons within the IC reduces the strength of ethanol-CTA in males, but not females, suggesting a sex-specific role for this neuronal subpopulation within the IC in mediating the aversive effects of ethanol.

Our finding that mIPSC frequency onto IC-BLA projections is higher following ethanol-CTA retrieval is consistent with our hypothesis founded on work showing increased mIPSC frequency onto L5/6 IC-BLA projections following LiCL CTA retrieval (Yiannakas et al., 2021), and suggests that this mechanism may be conserved across different taste aversion memories. Increased frequency of mIPSCs onto IC-BLA projections likely reflects stronger functional coupling between these cells and inhibitory interneurons. Indeed, the frequency of mIPSCs could be directly regulated by neurotransmitter release from PV interneurons onto IC-BLA projections. Such enhanced connectivity of inhibitory interneurons with principal cells is consistent with a critical role for interneurons in regulating excitatory and inhibitory balance within the IC to facilitate taste memory retrieval (Yiannakas et al., 2021). Although it remains to be tested, PV interneurons within the insula could also exert control over gamma oscillations, a role they play in other brain areas, including the hippocampus, to facilitate memory retrieval (Fuchs et al., 2007). Given that alterations in mIPSC frequency were limited to IC-BLA projections, rather than the general population of IC neurons, our findings suggest circuit specificity in plasticity resulting from the retrieval of ethanol-CTA memory. Furthermore, given that we observed no changes in mIPSC frequency or amplitude following a single, unpaired injection of ethanol, our results suggest that IC-BLA circuitry is responsive to the learned aversive association between ethanol and saccharin, and not ethanol alone. The finding that the effects of alcohol alone are dissociable from learning-induced plasticity resulting from ethanol-CTA is notable, and reinforces the strength of the CTA model in interrogating how alcohol’s aversive effects are encoded in the brain. Previous work suggests that the IC-BLA circuit is sensitive to abstinence from chronic alcohol exposure; withdrawal from 10, but not 7, days of chronic intermittent ethanol vapor increased AMPA-receptor-mediated postsynaptic function at IC-BLA synapses (McGinnis et al., 2020). That plasticity within this circuit is modified both acutely in response to a single-trial learned alcohol-associated aversion, and to longer-term withdrawal from chronic alcohol exposure, begs the question of how this circuit may shape alcohol consumption. In the present study, we observed a robust increase in inhibitory tone within this circuit 1 hour following retrieval of information about the aversive properties of alcohol, whereas previous work suggests a somewhat disparate effect; that is, 10 days of withdrawal from chronic alcohol exposure facilitated glutamatergic activity within the same circuit. One parsimonious explanation of these findings is that long-term exposure to alcohol disrupts the encoding of alcohol’s aversive effects via perturbation of excitatory and inhibitory balance within the IC, thereby disrupting the ability of IC-BLA projections to encode the aversive properties of alcohol. Future work should more directly consider how chronic alcohol consumption might dysregulate plasticity within the IC-BLA circuit to drive aversion-resistant drinking.

The finding that layer 2/3, but not layer 5/6 IC-BLA projecting neurons receive stronger local inhibitory input following ethanol-CTA retrieval is somewhat puzzling given increased frequency of mIPSCs following ethanol-CTA retrieval onto IC-BLA projections, regardless of layer. The lack of increased local inhibitory input to layer 5/6 pyramidal neurons, despite increased frequency of mIPSCs, could reflect differences in GABAergic inputs to these cells. One possibility is that GABAergic input to layer 2/3 IC originates locally, whereas GABAergic input to layer 5/6 arises from more distal projections and would thusly not be reflected in local synaptic input maps. Axons from the BLA preferentially and densely contact the more superficial layers of the IC, although the deeper layers of the IC are also innervated by the BLA to a lesser degree (Haley et al., 2016). Thus, another potential explanation for our observed increase in local inhibitory input to IC layer 2/3 following ethanol-CTA retrieval is a relative increase in BLA input strength to layer 2/3, over layer 5/6, IC interneurons, thereby driving L2/3 feedforward inhibition within the IC. This, coupled with the observation that the majority of uncaging-evoked events occurred in close proximity to the recorded cell’s soma, would be consistent with a role for layer 2/3 IC PV interneurons in regulating the function of local principal neurons.

Indeed, another key finding of the present study is that DREADD-mediated inhibition of PV interneurons within the IC reduced the strength of ethanol-CTA in males, but not females. Previous work suggests that PV interneurons are vulnerable to longer periods of alcohol exposure. For instance, six weeks of binge drinking, but not one week, increased the density of perineuronal nets surrounding PV interneurons within the insula, an alcohol-induced alteration that presumably facilitates inhibitory control onto local pyramidal neurons (Chen et al., 2015). Moreover, digestion of PNNs resulted in higher sensitivity to quinine-adulterated ethanol, thereby decreasing aversion-resistant drinking (Chen and Lasek, 2020). Our contribution to ongoing exploration into the role that IC PV interneurons play in mediating alcohol-associated behaviors suggests that these cells are responsive to alcohol in a more rapid timeline than has previously been uncovered. In combination with past literature, these findings indicate that across shorter periods of time IC PV interneurons are critical for an alcohol-associated aversive memory, but more chronically, upon repeated exposure to alcohol, may exert hyperactive inhibitory control over pyramidal cells in a manner that promotes aversion-resistant drinking. Of note, the effect of Gi DREADD-mediated inhibition of IC PV cells upon ethanol-CTA was modest; mice receiving a 2g/kg conditioning dose of ethanol in combination with DCZ-mediated activation of Gi-coupled signaling in PV interneurons still exhibited a robust, albeit blunted, taste aversion to saccharin during retrieval. One potential explanation for this small but significant reduction in CTA strength lies in the conditioning dose of ethanol used. Although there is precedent for a 2 mg/kg or higher dose of ethanol to elicit a robust ethanol-CTA (Barkley-Levenson et al., 2015; Cunningham, 2019; Gore-Langton et al., 2022), adult rodents routinely exhibit an ethanol-CTA to lower doses than the one used in the present study (Glover et al., 2016; Gore-Langton et al., 2022; Dornellas et al., 2024), suggesting the possibility of a near ceiling effect of our chosen dose in producing an aversion to saccharin. We also demonstrated a sex-specific role for PV interneurons in modulating ethanol-CTA, which was somewhat surprising given a lack of observed sex differences in GABAergic plasticity following ethanol-CTA retrieval. A 2g/kg dose of ethanol did not produce any observable differences in strength of ethanol-CTA between males and females in the present experiments; however, some evidence exists to suggest that females may be less sensitive to the aversive effects of ethanol relative to males across multiple behavioral paradigms, including ethanol-CTA (Sherrill et al., 2011; Morales et al., 2014) . A recent report noted that disruption of IC PNNs significantly increased sensitivity to quinine-adulterated ethanol in male mice, but not in female mice unless a very high concentration of quinine was added (Martins de Carvalho et al., 2023). Thus, the mechanisms driving both aversion-resistant drinking and strength of ethanol’s aversive effects may differ between males and females; our data lend support to the possibility that this difference may be driven in part by sex-dependent roles of IC PV interneurons in encoding ethanol’s aversive effects.

In summary, our data suggest that retrieval of an ethanol-CTA results in changes in GABAergic plasticity within an IC-BLA circuit, in a learning-dependent manner. Moreover, local inhibitory circuitry within the IC is significantly modulated by ethanol-CTA retrieval, and functional manipulation of PV interneurons within the IC suggests a sex-specific role for these cells in encoding the aversive properties of alcohol. These findings provide novel insight into circuit-specific physiological changes produced by a single associative learning experience regarding alcohol’s aversive properties. Future work should examine how long-term ethanol consumption might alter excitatory and inhibitory plasticity within the same IC-BLA circuit to mediate aversion-sensitive drinking.

## Acknowledgments

This work was supported by the National Institutes of Health (NIH) National Institute of Alcohol Abuse and Alcoholism (NIAAA) grants T32AA007573 and F32AA031395 to LRT, F32AA030494 to HLH, K99AA030628 to MEF, and P60AA011605 to TLK and the Bowles Center for Alcohol Studies

